# Neural correlates of rapid familiarization to novel taste

**DOI:** 10.1101/2024.05.08.593234

**Authors:** Daniel A. Svedberg, Donald B. Katz

## Abstract

The gustatory cortex (GC) plays a pivotal role in taste perception, with neural ensemble responses reflecting taste quality and influencing behavior. Recent work, however, has shown that GC taste responses change across sessions of novel taste exposure in taste-naïve rats. Here, we use single-trial analyses to explore changes in the cortical taste-code on the scale of individual trials. Contrary to the traditional view of taste perception as innate, our findings suggest rapid, experience-dependent changes in GC responses during initial taste exposure trials. Specifically, we find that early responses to novel taste are less “stereotyped” and encode taste identity less reliably compared to later responses. These changes underscore the dynamic nature of sensory processing and provides novel insights into the real-time dynamics of sensory processing across novel-taste familiarization.

## Introduction

Gustatory cortex (GC) is the “primary” sensory cortical taste area, wherein neural ensemble taste responses both code taste quality and play important roles in driving gustatory perception. Specifically, GC cortical responses progress through a sequence of metastable states (Jones et al., 2007; La Camera et al., 2019) the last of which is causally related to the onset of taste-consumption behavior (Sadacca et al., 2016; Mukherjee et al., 2019). While the speed with which this dynamic unfolds differs from trial to trial, the ensemble-dynamic itself is reliable enough that it can be assessed in single trials, such that the onset latency of the “late” state following delivery of an individual aliquot of bitter taste can be shown to directly precede the rejection of that aliquot.

This single-trial clarity of response offers an unprecedented opportunity to test the extent to which the cortical taste-code is innate versus “learned.” The “standard” model of taste perception holds that taste is an “innate” and “predetermined” sense; it has been suggested that learning is not required to generate appetitive and aversive responses to taste because these responses are generated by “hardwired” taste circuits. The “elegant simplicity” of such a system is appealing because it’s difficult to imagine how codes that change and drift over time could generate consistent and robust behavioral responses. There is, however, a good deal of evidence that cortical sensory representations do change over days: exposure to tastes, for instance, changes taste responses in GC between sessions (Flores et al., 2022; Staszko et al., 2022); demonstrations of neural drift (Schoonover et al., 2021) similarly suggest a certain lack of sensory-circuit hard-wiring. Given these findings, and the fact that lab rodents are nearly 100% “taste naïve” when experiments begin, it is reasonable to ask whether such “refinements” might be observable across smaller time scales—whether responses change even across the initial trials of novel taste exposure. This question can be answered by making use of the single-trial characterizability of GC taste responses.

Here we do so, bringing analyses used to capture dynamic ensemble-level characterizations of taste responses in single trials to study how these responses evolve across initial trials of taste exposure. These analyses reveal that responses from the earliest ∼10 trials of taste exposure cluster distinctly from those from later trials; this separation into early- and late-session trial types is most reliable in truly taste-naïve rats, suggesting that these rapid changes are likely “experience-dependent”(as opposed to randomly drifting), but can also be observed to a lesser degree in 2^nd^ taste exposure sessions, suggesting the gradual learning of an “experimental set.” Examined separately, the initial taste responses were found to contain all of the hallmark temporal properties of “normal” taste responses. Furthermore, the ensemble states of those initial responses code taste-identity less reliably than those of later responses.

Overall, this work adds substantial insight to the increasing literature describing experience-dependent changes in sensory processing, by presenting what is to our knowledge the first evidence that a significant amount of change in sensory processing occurs with initial exposure trials, in real-time as animals receive novel sensory experience, and that these changes are likely tied to experience-correlated changes in consumption behavior.

## Methods

### Experimental design

#### Subjects

Female Long–Evans rats (*n* = 8); 250–300g prior to surgery) were used for this study. Rats were maintained on a 12 h light/dark schedule and were given *ad libitum* access to food (Labdiet 5P00 Prolab RMH3000) and water before experimentation. To minimize taste preexposure, we ensured rats housed singly, in clear plastic cages containing only sawdust (or paper post-surgery) bedding, and paper enrichment. All use of animals followed the Brandeis University Institutional Animal Care and Use Committee guidelines.

#### Surgery

Intraperitoneal injection of a ketamine/xylazine mixture was used to anesthetize the rats (100, 5.2, and 1 mg/kg, respectively). Supplemental intraperitoneal injections were given as needed. Once under anesthesia, the rat was positioned within a conventional stereotaxic apparatus, and its scalp was removed. 5 small burr holes were drilled into the skull for the placement of 0–80 ground screws. 2 larger craniotomies and durotomies (∼2mm dia.) were made in the skull, dorsal of GC. An electrode microdrive (see *Electrophysiology*) was implanted bilaterally into the gustatory cortices, and electrodes were positioned 0.5 mm above the GC (+1.3mm A/P, +/- 5mm M/L, 4.6mm D/V). Afterwards, craniotomies were sealed with Kwik-Sil (World Precision Instruments; Sarasota, FL), the microdrive and one intraoral cannula (IOCs; referenced in Katz et al., 2001) were secured to the skull using dental acrylic.

#### Passive taste administration paradigm

Three days following surgery, electrode bundles in the microdrive were driven down 0.5mm to the depth of GC, and animals were placed on a water-restriction regimen (30mL water/d), then habituated to the experimental environment for 2 d, followed by 2d habituation to 30μl water deliveries through the IOCs. Following habituation sessions, animals were exposed to the experimental taste-solution battery [0.3M Sucrose (Suc), 0.1M Sodium Chloride (NaCl), 0.1M Citric Acid (CA), and 0.01M Quinine Hydrochloride (QHCl)] through a manifold of fine polyimide tubes inserted to 0.5 mm past the end of one IOC (eliminating any chance of mixing) and locked onto the dental acrylic cap. All fluids were delivered under ∼20PSI of nitrogen pressure; while delivering each taste from one side may have meant not entirely immediate exposure of all taste buds, the pressure ensured that the brief (∼40-100ms) taste administration onto the tongue was complete before any taste-related dynamics appeared in GC responses) resulted in extensive tongue coverage at reliably short latency (Katz et al., 2001), and delivery through a single manifold ensured consistent presentation of all stimuli. In each taste-administration experiment, after rats were situated in the experimental arena, they were allowed to sit in the inert arena for 20 minutes prior to the start of taste exposures. Rats then received 30 deliveries of each taste (120 trials total) in a pseudorandom random order, separated by randomly varied inter-trial intervals of 20-30s each. Total fluid delivered was 3.6 ml per session over 40-50 minutes. After removal from the experimental arena, animals were given access to 30mL water. The experimental environment wherein animals were subjected to the paradigm consisted of a small, fully enclosed acrylic 1×1×2-foot arena, encapsulated by a larger opaque-and-faraday-caged box designed to insulate the animal and arena from light as well as auditory and electronic noise.

Animals were subjected to 3 consecutive sessions of taste exposure. In some cases, we were only able to record 2 consecutive sessions of taste exposure.

#### Electrophysiology

Neural probes implanted into rats consisted of two independently drivable electrode bundles separated by 10mm (the distance between the gustatory cortices), each consisting of 32 nichrome microwires (64 total), that were affixed to an electronic interface board (EIB) (Open-Ephys). Throughout each taste-exposure session, neural signals were simultaneously amplified and digitized by a 64-channel headstage (Intan, Open-Ephys) connected to the EIB, and recorded at 30kHz using an Intan RHD USB interface board (catalog# C3100) for offline analysis. In addition, timestamps of each stimulus delivery were recorded alongside electrophysiology.

#### Histology

Prior to surgery, electrode tips were coated with Vybrant Dil cell-labeling solution (Invitrogen, catalog #V22885) to label the cells adjacent to the electrodes with red fluorescent dye. In preparation for histology, rats were deeply anesthetized with a ketamine/xylazine mixture, and centrally perfused using 0.9% saline followed by 10% formalin. The brain was then harvested and incubated in a fixing mixture of 30% sucrose and 10% formalin for 7d. Brains were sectioned on a sliding microtome (Leica SM2010R, Leica Microsystems) into 50 μm coronal slices. Sections contaiing the electrode sites were then mounted on histology slides and labeled with a green fluorescent Nissl stain (Invitrogen NeuroTrace, catalog# N21480). Slides were then imaged at 2x using an automated fluorescent microscope (Keyence BZ-X800), and the location of GC was validated by comparing the anatomical features of the imaged sites containing electrode tracks with a rat brain atlas (Allen Brain)

### Data Analysis

#### Electrophysiology Data Preprocessing

Electrophysiology was loaded into the blechpy analysis package (Nanu and Svedberg, 2023), where candidate spikes were identified as signals with at least a 3:1 signal-to-noise ratio. Candidate spikes were then clustered and sorted to identify putative spikes: the UMAP algorithm was used to extract waveform features from candidate spikes, which were fit by Gaussian mixture models to cluster spiking events. Clusters were then manually curated. Only clusters clearly separated in UMAP or PCA space, with waveforms that appear visually consistent with one another, containing zero 1ms inter-spike-interval (ISI) violations and fewer than 1% 2ms ISI violations, were chosen as putative spikes. Visual inspection of waveforms was performed to putatively determine if the unit was a pyramidal neuron or fast-spiking neuron. Spikes identified by this process were then drawn into peri-stimulus spike arrays for each trial according to the timestamps demarcating the onset and identity of each stimulus. These methods and criteria are consistent with previously published methods from our lab (Moran & Katz, 2014; Sadacca et al., 2016; Levitan et al., 2019; Mukherjee et al., 2019).

#### Analysis of single-unit responses

Firing rates for each non-overlapping 250ms bin were calculated to compute firing-rate arrays for each trial, which were averaged across trials to create the peri-stimulus time histogram (PSTH). To calculate taste-discriminability, a one-way ANOVA of the firing rate in each time-bin was computed across taste stimuli for responses in the subset of neurons that were held across at least 1 pair of days. To compute correlation with palatability, the Spearman correlation coefficient (R2) was calculated between the firing rate and the “canonical” palatability rank of the taste stimuli, for each bin, for each taste responsive neuron. In both the analyses of taste discriminability and palatability correlation, sequences of three consecutive bins with a p-value < 0.05 were marked as significant. Population-level palatability-correlation was calculated as the average palatability-correlation in each time bin, averaged across all recorded neurons that exhibited significant palatability correlation in their responses.

#### Poisson Hidden Markov Models (HMMs)

To extract the temporal structure of ensemble activity on the scale of single-trials, we fit Poisson HMMs to spike trains from simultaneously-recorded units (Jones et al., 2007; Moran and Katz, 2014; Nanu et al., 2021). We fit separate models for each tastant for each recording session from 0ms to 2000 ms peri-stimulus, with 2-6 states connected in a feed-forward manner. The input model was fed with neuronal spike data at a resolution of 1bin/ms, and the initial spike frequencies for each neuron were determined based on a normal distribution reflecting the average and variability of their firing rates over multiple trials. Starting values for the transition matrix were also chosen randomly from a normal distribution, with a set mean and a low variance, to favor stability in state retention over time steps. The main diagonal of this matrix, which reflects the likelihood of maintaining the current state, was set according to a distribution emphasizing high state retention, while the probabilities of reversing or skipping states were set to null. All transition probabilities were then adjusted so that each row summed to 1. The models were individually fit to each dataset 50 times using the Baum-Welch algorithm, with the fitting process concluding once the change in the model’s log likelihood between updates fell below a specified threshold. Across iterations for each parameter-set that converged above the log-likelihood threshold, the rendition with the best Bayesian Information Criterion (BIC) was considered the most accurate for the data. To determine which number of states best-fit the data, we chose the model that minimized the Akaike Information Criterion (AIC) for each modeled set of trials, allowing the number of states to vary between trials, in order to be able to analyze changes in coding that occurred across trials in a session. State sequences for each trial determined by calculating the state with the highest emission-probability in that time-bin.

#### Calculation of trial-distance matrices

Distance matrices for each “set” of trials—the responses to a specific taste in a specific recording—were produced by the following procedure: Euclidean distance of each pair of trials was calculated as the distance of the firing-rate-array vectors (see Analysis of single units) for each matching time-bin from 100-2000ms post-stimulus. These trial-distance vectors were then averaged to calculate the average distance between the pair of trials.

#### Calculation of consensus clustering

Each trial-set’s distance matrix was subjected to hierarchical clustering using the scipy.cluster.heirarchy functions from the Scipy python module (SciPy, 2022). The silhouette score method was used to tune the threshold of each clustering, using the sklearn.metrics.silhouette_score functions from the scikit-learn Python module (scikit-learn, 2022). These data were then aggregated using a modified consensus clustering algorithm (Monti et al., 2003) to find the consensus distance matrix. Briefly: the number of trial-sets in which each pair of trials were not given the same cluster-label were counted and divided by the total number of trial-sets to produce the consensus-distance for each pair of trials. These consensus distances were then plotted as a correlogram using the imshow function from the Matplotlib Python module (Matplotlib, 2024). Heirarchical clustering of the correlogram was also plotted as a dendrogram using the SciPy dendrogram function.

#### Analyses of hierarchical clustering

To identify the number of trials in- and average trial number of-the largest two clusters, the trial numbers and cluster labels belonging to the two largest clusters in each trial-set’s hierarchical clustering of the distance were identified. For the 2 largest clusters of each trial-set, the one with the lowest average trial number was identified as the “early” cluster, and the one with the highest was identified as the “late” cluster. To create a null-distribution of the clustering, 100 iterations of the following was performed: each the trial-numbers of each trial-set were resampled with replacement, and the above analyses were re-performed for each iteration. Significant differences were calculated using a two-tailed permutation test to compare the sample means of the % of trials in cluster and average trial number to the corresponding null-distribution, where sample means exceeding the 97.5^th^ percentile or falling below the 2.5^th^ percentile of the null-distribution are considered significant with a p-value < 0.05.

#### Euclidean distance to template (EDT)

For each set of trials considered within a template, the “template trial” was calculated as the trial-averaged firing-rate-vector over each ensemble was calculated for each time-bin 100-2000ms post-stimulus. The Euclidean distance between the firing-rate vectors of the sample-trial and the template-trial were then calculated for each bin and averaged across bins to calculate each-trial’s EDT. For analyses performed on this measure across many sets of trials, the EDT of each trial was z-scored across each set of trials coming from an ensemble, to normalize Euclidean distances so that they can be compared across ensembles of different sizes, since larger ensembles naturally produce longer distances due to higher dimensionality.

#### Split-session analysis of EDT

For each trial set, trials were bisected across every possible trial, and for every bisection point, the EDT was calculated for each sub-set of trials on either side of the bisection point. The “best” bisection was simply identified as the one with the lowest average EDT across trial-sets. Statistical comparison of “trial-split” EDT against single-template EDT was done by performing a repeated-measures, within-subjects ANOVA using the pingouin Python module (Vallat, 2024), comparing the EDTs of the “best” trial-split against the single-template EDTs for each trial-set, using animal ID as the subject variable and the repeated-measures variables were taste and “trial-split” versus “single-template”. Similarly, statistical comparison of EDT for “early-session” and “late-session” trials within the best trial-bisection was performed as a repeated-measures within-subjects ANOVA, where animal ID was the subject variable and taste and “early-session” and “late-session” were the repeated-measures.

#### Logistic Function modeling

We used a modified exponential decay function to model the relationship between various variables and trials: 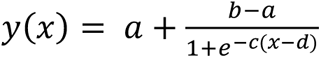, where trial-number is x. Coefficients a,b,c and d were fit using the scipy.optimize.curve_fit function from the SciPy Python module (SciPy, 2024). We placed the following constraints on these coefficients: c must be positive, a and b must not exceed the bounds of the data, and C cannot be larger than the range of the data divided by the number of trials. This function was chosen because it naturally achieves the nonlinear-changes we hypothesized characterizes our data, and because it is easily constrained for curve fitting algorithms. For our analyses of EDT, transition-time, and state-accuracy, the independent variable was modeled for each set of trials of a specific taste in a specific recording. Once each trial-set was modeled, the average-model was calculated by bootstrapping the mean value, as well as the 95% confidence interval, of the independent variable across all the models over 10,000 iterations, for each time-step. The coefficient of determination (R^2^ value) of each model was calculated as one minus the sum of squared residuals to the model divided by the sum of squared residuals to the empirical average of the independent variable. The “predicted change” of each model was calculated as the predicted value of the independent variable for the last trial minus the predicted value for the first trial. The average R^2^ and predicted change are calculated as the empirical mean across trial-sets.

#### Trial-shuffled logistic function modeling and permutation tests

To control for overfitting, we compared the analyses of logistic function modeling to equivalent analyses of trial-shuffled data, where shuffling was performed across 10,000 iterations. For each iteration of shuffling, the trial-number of each trial-set’s data was simply shuffled without replacement, and the same analyses performed on the sample-data were performed on each iteration of the shuffle-data. Averages of shuffle-data statistics were then taken grouping by iteration-number to create null-distributions of the R^2^ values and predicted change. These null-distributions were then used to perform two-tailed permutation tests of the R^2^ and predicted-change sample-means, where sample-means falling outside the 97.5% confidence interval (Bonferroni corrected for the number of comparisons) of the null-distribution are considered significant with a p-value < 0.05

#### Bayesian decoding of taste identity from HMM states

Because for computational-efficiency reasons, the HMM was only performed on trial-sets of responses to each specific taste in each recording. Therefore, in order to compare the coding of HMM states across HMM solutions for different tastes in the same recording, activity (firing-rate-vectors across trials) from each state across all the HMMs in a recording was used to train a gaussian naïve-Bayesian classifier (SciKit-Learn, 2024), where each state was identified as the combination of it’s taste and state-number (i.e. “Suc_1”, “NaCl_3”, etc.—if a recording contains 4 HMM states per taste and 4 tastes were given, the classifier would decode from 16 possible states). Each HMM-state in each trial was then re-classified as the probability that the activity would be decoded as the state it came from (as well as the probability of every other possible state), using the jackknife method for cross-validation. Probably of decoding the correct taste was then calculated as the sum-probability of decoding any one of the states for the same taste.

#### Selection of “Early” and “Late” states

Previous work—on the assumption that the identity and latency of HMM states would be relatively consistent across trials—selected “early” and “late” trials by finding the longest states within certain time-periods post-stimulus (Sadacca et al., 2016). Since these analyses specifically would interrogate changes in the temporal dynamics and identity of these states, we devised a criterion that could find “early” and “late” states that change in identity across trials or dramatically shift in latency to onset from trial to trial. This involved selecting the two states in each trial that—met the criteria of starting at least 50ms post-stimulus and having a duration of at least 50ms—with the highest probability of decoding the correct taste in our Bayesian analysis. Of these two states, the one that emerges earliest in the trial is considered the “early” state and the one that emerges later is considered the “late” state.

## Results

### Recording of GC taste responses reveal the expected dynamics across trials and sessions of novel taste exposure

We bilaterally implanted taste naïve rats with electrodes in gustatory cortex (GC) and exposed them to a battery of taste solutions—30 trials each of 0.3M sucrose (sweet), a palatable low concentration—0.1M—of sodium chloride (salty), 0.1M citric acid (sour), and 0.001M quinine hydrochloride (bitter), randomly ordered, for 3 consecutive days. We then analyzed these data to determine changes in the dynamics of taste responses from single neurons and ensembles.

We validated that we could record and isolate single neuron ensembles (Fig 1A), that many of these neurons exhibit the previously-described dynamics of taste-responses (Fig 1B), significantly discriminating taste across all three sessions (Fig 1C). As expected (Sadacca et al., 2016), palatability-related firing peaked at the beginning of the 2^nd^ second of the responses (Fig 1D).

**Fig. 1:**
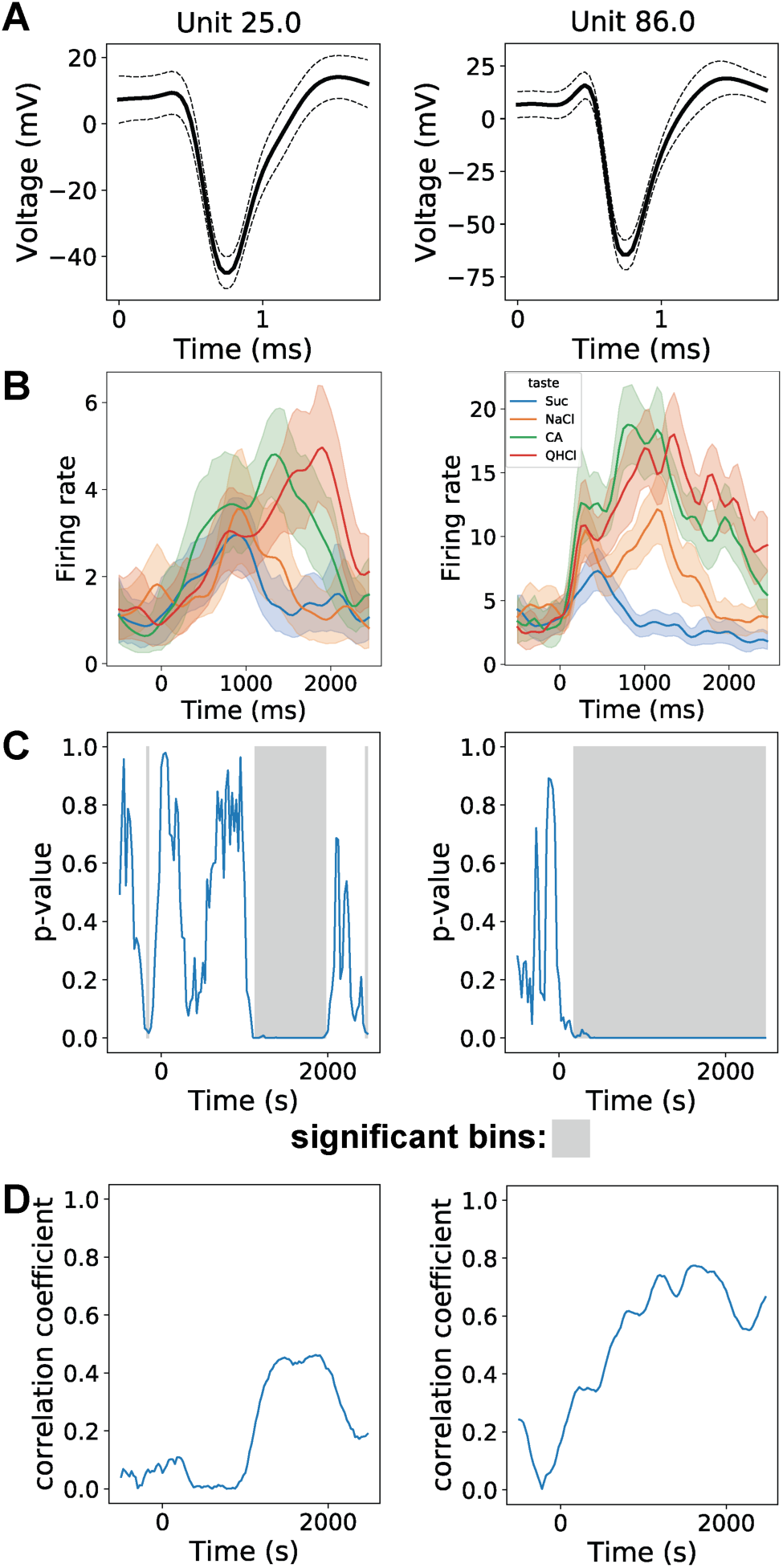
Cortical responses to novel taste exposure are taste-specific and palatability-related. **(A)** Waveforms of 2 representative GC single neurons held across sessions of novel taste exposure. **(B)** Peri-stimulus-time-histograms of taste-responses to Sucrose (Suc), Sodium Chloride (NaCl), Citric Acid (CA), and Quinine Hydrochloride (QHCl) from held units in (A). **(C)** Quantification of taste discriminability of responses in (B). Gray region indicates period when taste is significantly (p < 0.05) discriminable (p < 0.05). **(D)** Quantification of taste-responses’ (Spearman) correlation with palatability-rank, showing that the palatability-relatedness of GC taste responding peaks late.

Studying changes in taste responses the scale of individual trials requires parsing the ensemble-level temporal dynamics that define them. We used Hidden Markov Modeling (HMM) to extract ensemble-level state-transitions from population activity in sets of responses to each taste (Fig. 2A). An advantage of this approach is that the rapid transitions in ensemble-level activity are captured, which would otherwise be smoothed over by any measure that averages over many trials (Fig. 2B).

**Fig. 2:**
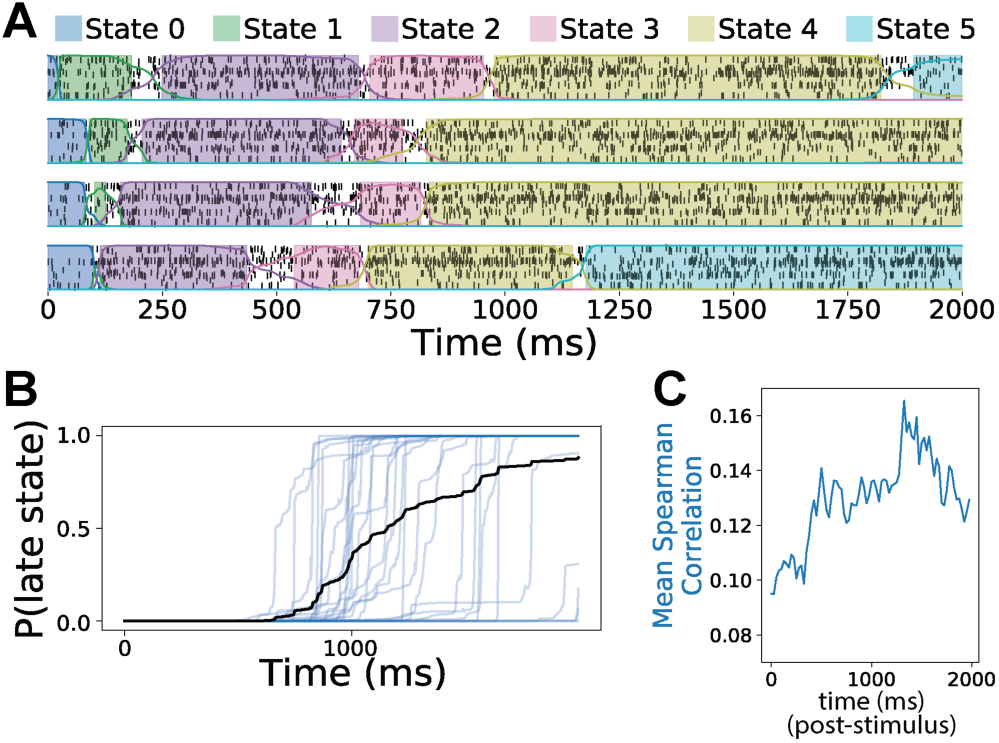
Novel taste exposure elicits reliable sequences of sharp state-transitions from cortical ensemble activity. **(A)** Example of state transitions revealed by Hidden Markov Modeling (HMM) of cortical ensemble activity in response to 4 trials of taste (sucrose) stimulation. Spikes of every neuron in the ensemble are marked vertical black hash marks. Colored lines indicate probability of each state, and regions billed by matching colors indicate times where probability of the state is greater than 0.8. **(B)** Representative example of a session of transitions from the early to late epoch for the same animal depicted in (A). Blue lines depict individual transitions, and the black line depicts the average of the transitions, revealing that state-transitions in a single trial are sharper than they are across a session. (**C**) Quantification of the average correlation with palatability rank across all recorded neurons in this example, showing that palatability rises as the probably of the late state rises in (B).

### Taste responses across trials within a session organize into two clusters

Multiple studies on this system, from this and other labs, (Sadacca et al., 2016; Mukherjee et al., 2019; Arieli et al., 2022; Lang et al., 2023)—as well as in other systems—have extensively documented what is also obvious in Fig 2B—the trial-to-trial variability in the timing of reliably appearing dynamic features of taste responses. These studies have assumed that this variability is random—that trials at all points within a session are equally likely to reflect the average trial. Underlying this assumption is another: that rats arrive at this tasting session ready to maturely process tastes.

Given the many studies documenting the effect of familiarization on taste perception and coding between (Flores et al., 2022; Staszko et al., 2022; Schiff et al., 2023) and even across (Monk et al., 2014) sessions, it is reasonable to question this assumption, and to ask whether the initial trials of taste exposure might engender response dynamics that are distinct from those observed in the “mature” average of trials. We therefore asked whether there might be differences in information content coded in states within early taste responses (compared to later responses), or significant differences in the latencies of these states’ onset, examining the stability of taste responses in a manner that is agnostic to the within-trial temporal dynamics, by analyzing the firing rates of neurons in each ensemble across a stationary window of time post-stimulus (100-2000ms), a window of time that encapsulates the majority of taste-processing. We reasoned that if coding changes systematically across trials, then trials closest together in time should bear the most similar taste-responses. Conversely, if taste-responses occupied a single, stable regime throughout the session, then variability across trials should be unimodally distributed around this singular state, and the dispersion of this variability should be unrelated to trial number.

To test this null hypothesis, we calculated the Euclidean distance between each pair of trials (using ensemble firing-rate vectors) in each session of responses to a particular taste, to produce a trial-distance matrix and subjected to hierarchical clustering, which was then used to produce a consensus-clustering distance matrix (Fig 3A) in which each node represents the frequency that a pair of trials were not clustered together. Hierarchical clustering of the consensus distance matrix—represented as a dendrogram—was performed to illustrate how trials organized into groups (Fig 3B).

**Fig. 3:**
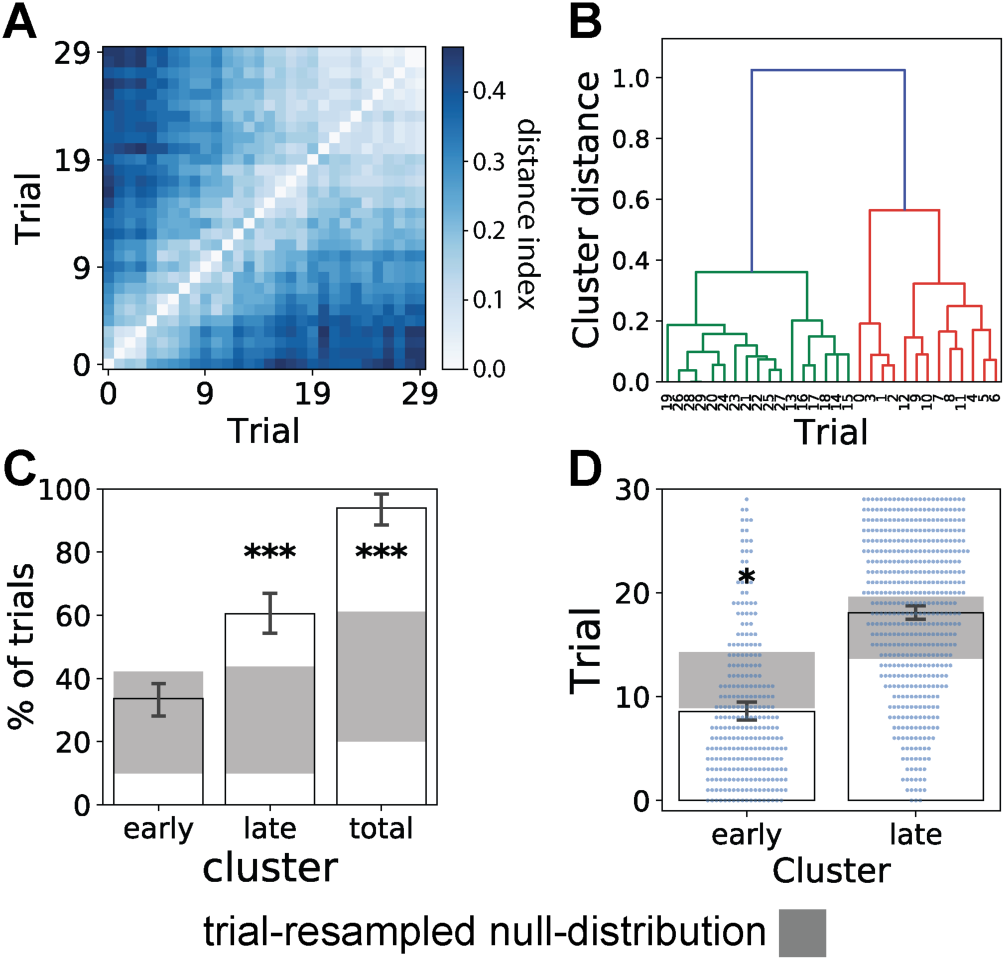
GC ensemble taste responses reliably form 2 clusters separating early- and late-session trials across novel taste exposures. **(A)** Distance matrix of hierarchical consensus clustering of population-firing-rate-vectors for each pair of trials, across each set of exposures to a taste in the first session of taste exposures, for each animal. **(B)** A dendrogram of consensus clustering in (A) reveals trials fall into two distinct clusters. **(C)** Quantification of the % of trials that fall into the two largest clusters, for each set of exposures to each taste, for each animal. As a control, a null distribution was created by randomly resampling trials with replacement and repeating the clustering procedure on the resampled trials. The 97.5% confidence interval) of the null distribution is plotted (gray). In each session, significantly more trials were clustered into the two largest clusters than would occur to chance alone (p < 0.001, permutation test, Bonfferoni corrected for 9 comparisons). **(D)** Quantification of average trial number in each cluster (bars graph, +/- 95% confidence interval) versus trial-shuffled control (97.5% confidence interval, gray box). Each individual trial belonging to the “early” and “late” clusters are depicted as blue dots. Average trial number of the “early” cluster is significantly lower than predicted by chance.

These analyses suggest that trials tend to organize into two distinct clusters made up of trials that occur close to one another in time, and that more specifically the earliest trials cluster distinctly from those making up the bulk and balance of the session. In order to rigorously test these appearances (that trials primarily organized into two clusters and that these clusters significantly separated early-session and late-session trials, we first quantified the percentage of trials in the two largest clusters for each individual trial-set, and compared this to the percentage of trials that would organize into the two largest clusters by chance—calculated by computing a null-distribution where trials were resampled with replacement (Fig 3C). Reliably, the percentage of trials that organized into the two largest clusters exceeded 90%—significantly more than would occur by chance alone (p < 0.001). Next, we tested if the trial-number of the two largest clusters is significantly polarized (as opposed to randomly dispersed between them). For each clustering, the average trial number of the two largest clusters was calculated to determine which was the “early” cluster and which was the “late” cluster. The average trial-number of the early and late clusters was then compared to the corresponding null-distribution calculated in the same way from the trial-resampled data, to determine if the dispersion of trials between the early and late clusters is significantly more polarized than chance (Fig 3D). This analysis reveals that only the early cluster in the first session had a significantly lower average trial-number than chance (p < 0.05), revealing that the dynamics of taste responses change early in the session.

### Taste responses become more stereotyped across trials of novel taste exposure

On the basis of the above findings, we hypothesized that GC ensemble taste responses “settle into” the stereotypical patterns observed in many previous papers across the first portion of the session. To test this implication—that a session of novel taste exposure can be characterized as showing a “regime-change” that bisects the session into a set of early trials and a set of later trials that induce more “mature” taste response dynamics—we first reasoned that separate templates constructed from the averages of earlier and later trials should better represent the entire session of taste responses than an overall average-constructed template. We performed this analysis for every possible bisection point, calculating average (Z-scored) distances between each template and each trial used in its construction. This revealed that for each session, the best-division converged around a specific trial that produces the two “optimum” templates, where the average distance of each trial to its template is minimized when trials 0-13 and 14-29 are considered separately (Fig 4A), and that the optimized two-template model significantly better-represents the trials compared to a single-template model (the average Euclidean distance of each trial to the average response for all trials) by around 0.4 standard deviations (p < 0.001) (Fig 4B)—a finding consistent with the idea that taste responses switch from one regime to another over the course of taste exposures.

**Fig. 4:**
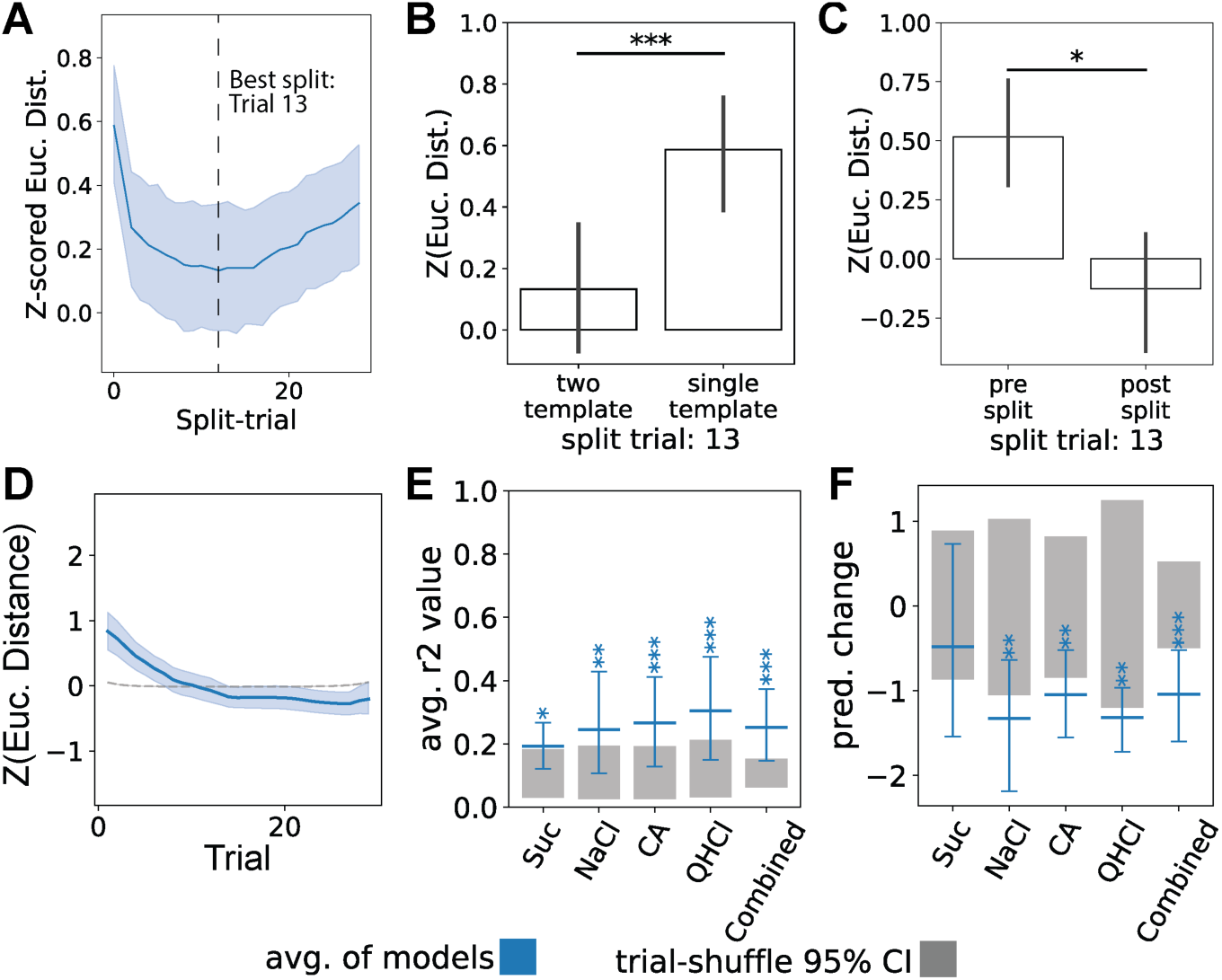
Clustering of GC ensemble taste trials reflects the process of responses become stereotyped. **(A)** Each session was repeatedly bisected into two parts with a sliding bisection point, and average responses (templates) were made of each pair of subsets (see text). Presented are Z-scored Euclidean distances to the corresponding templates for each bisection of trials, averaged across all trials (blue curve), with 95% confidence interval (shaded interval). Lower distances therefore connote better fits to the templates. The best fits are crowded to the left, demonstrating—consistent with Figure 3—that the best bisections divide the few early trials from the bulk of the later trials. **(B)** The average Z-scored Euclidean distances of each trial to the corresponding template, for the case in which trials are divided at trial 13, is compared against the single-template case, revealing that the two-template case significantly better-represents trials than the single-template case. **(C)** The average Z-scored Euclidean distances of each trial to its corresponding template, when trials are divided at trial 13, reveals that the post-split trials are significantly more similar to one another than the pre-split trials (*: p < 0.05, **: p < 0.1, *** : p < 0.001). **(D)** The Euclidean distances of each trial’s taste responses (as a population firing-rate vector) from the “stereotyped” response, modeled using a logistic function for each set of responses pertaining to each taste, session, and animal, and then bootstrap-averaged for each trial (blue line). Trial-shuffle/control models (gray dashed line) were generated *via* 10,000 iterations of trial shuffling with the modeling procedure repeated for each shuffle. **(E)** The average coefficient of determination across logistic models, +/- the 97.5% confidence interval is plotted in blue, and compared to the 97.5% confidence interval of the trial-shuffle null-distribution (gray), revealing that on average, stereotypy increases with trial-number, crossing the control line at around trial 10. **(F)** The average across-session change (last trial – first trial) predicted by the models plotted in (A) is plotted as blue horizontal lines; error bars represent the 97.5% confidence interval of the mean (determined *via* bootstrapping). These are plotted against the 97.5% confidence interval of the corresponding null-distributions (gray box) calculated from the trial-shuffles. * = 0.05 ; ** = 0.01; *** = 0.001.

Next, we tested whether these two taste-processing regimes seen across a session of novel taste exposure are best described as occupying two stable states, or rather (as we hypothesized) as an unstable state (a "transient,” perhaps) giving way to a stable state. To do this, we compared the ability of early- and late-trial templates to predict each trial in its regime (Fig 4C), which revealed that the responses falling into the early group of trials were significantly further from their template on average than those in the late group of trials (p < 0.05). Thus, the trial-to-trial variation across novel taste exposures is best described as an unstable regime that gives way to a more stable one. Under the context of the broader hypothesis that changes in taste responses are experience-driven, it is reasonable to suggest this could mean that taste-responses progressively converge upon a stable regime of coding over the course of taste exposures.

‘Convergent evidence for our conclusion that novel taste responses progressively “settle into” a stable regime came from an examination of the relationship between the (Z-scored) distance of each taste-response and the full-session template across trial number. When we modeled this relationship using a logistic function, we observed a clear converge to a stable mean (Fig 4D). By trial 13, responses had stabilized. Furthermore, the average coefficient-of-determination (R^2^) value of the logistic models applied to the actual session data was significantly higher than a null-distribution of fits computed by repeating the distance calculation and modeling of distance-to-template on trial-shuffled data over 10,000 iterations (Fig 4E); this analysis revealed that increases in trial stereotypy are correlated with trial number, both for all tastes (p < 0.001) and when models are averaged separately for each taste (Suc: p < 0.05, NaCl: p < 0.01, CA: p < 0.001, QHCl: p < 0.001).

Finally, we performed a direct test of whether taste responses converge to a stable regime. We reasoned that if so, the models should demonstrate that the distance to the template-response consistently decreases across trials. To test this prediction, we quantified the average magnitude of change in stereotypy predicted across the models and compared this to the null-distribution control (Fig 4F). As predicted, the Euclidean distance to the template significantly decreased across the span of a session (p < 0.001), indicating that taste-responses become substantially more stereotyped with taste exposure; this was true for the overall dataset, and for 3 out of the 4 tastes tested individually. Together, these results allow us to confidently conclude that taste responses converge onto a stable regime, suggesting a “refinement” of taste responses across novel taste exposures.

### Analysis of dynamic states reveals that changes in coding, not temporal dynamics, drives refinement of taste responses across novel-taste-exposure

Since taste responses are characterized by temporally dynamic sequences of metastable states (Jones et al., 2007; La Camera et al., 2019), analysis of these dynamics is necessary if we are to dissociate whether: 1) changes in GC responses across exposures are driven by changes in the information coded by the ensembles themselves; or 2) these early trials contain otherwise-normal taste responses emerging with abnormal temporal dynamics. We therefore performed Hidden Markov Modeling (HMM) on each set of responses to each taste in each session, thereby qualitatively revealing both shifts in the state-identities that appear in taste responses and in these responses’ temporal dynamics (note that the early trials shown at the top of Fig 5A look very different from later responses, both with regard to the nature of dominant states and the times of state transitions).

**Fig. 5:**
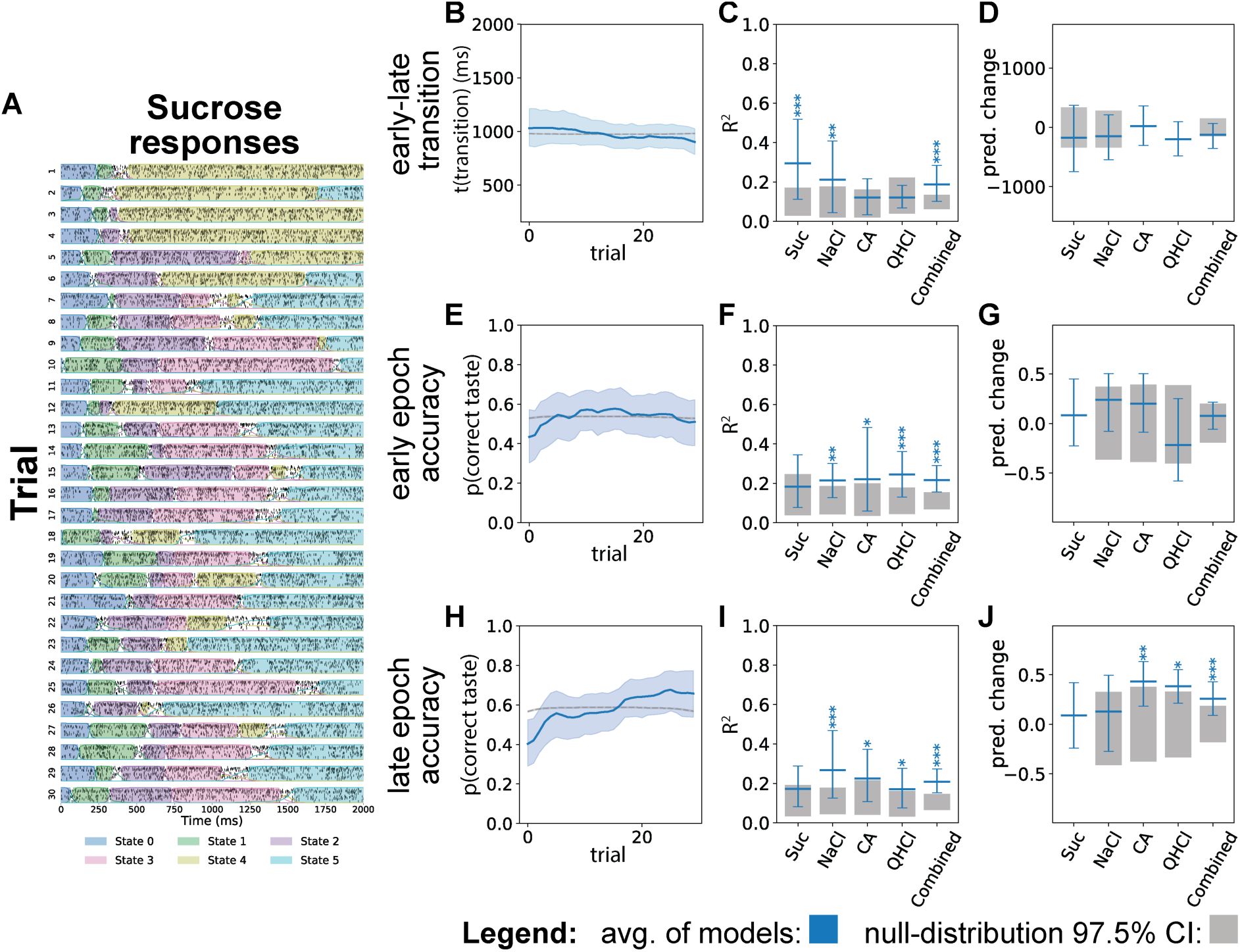
The timing and reliable taste-specificity of ensemble state transitions change across trials of novel taste exposure. **(A)** A representative session of HMM sequences across exposures to sucrose, reveals a qualitative change in patterning. **(B)** Early-to-late epoch transition latencies for each trial across the session +/- 95% confidence interval (in blue), related to the averaged trial-shuffled equivalent (dashed gray line). **(C)** Average coefficients of determination (R^2^) for the models of (B) reveals a significant association between trial-number and transition time—while the across-session change is small (see D), the non-flat nature of the function is reliable. **(D)** Quantification of the change predicted in (B) reveals a near-significant change from 1^st^ to last trials in the latency of the early-to-late transition. **(E)** Taste identity-coding accuracy in the early-epoch for each trial across sessions (generated as in E). **(F)** Average coefficient of determination (R^2^) for the models in (E) reveals an association between trial-number and early-epoch taste-coding accuracy—coding became more reliable after the first few trials. **(G)** Quantification of the change predicted in (E) reveals that taste-coding accuracy of the early epoch does not reach significance. **(H)** Taste-identity-coding accuracy in the late-epoch for each trial across sessions of novel taste exposure (generated as for C). **(I)** Average coefficient of determination (R^2^) for the models in (H) reveals an association between trial-number and late-epoch taste-coding accuracy. **(J)** Quantification of the change predicted in (I) reveals that taste-coding accuracy of the late-epoch significantly increases across trials of taste-exposure (p < 0.001).

The most salient feature of the dynamics that can be observed in HMM analysis of taste responses (see late trials in Fig 5A) is the transition from the early “identity-coding” state that dominates the first ∼100-700ms post-stimulus, to the late “palatability-correlated” state that dominates the activity following the early state (Sadacca et al., 2016). Previous work has shown that this transition is causally related to the onset of aversive consummatory behaviors (Mukherjee et al., 2019), making the timing of this specific transition of particular interest. To test the hypothesis that the early-to-late state transition changes across novel taste exposures, we first modeled the relationship between trial number and the time of this transition across each set of trials using a logistic function (Fig 5B). While visual inspection of this relationship suggests that trial-to-trial differences are small, analysis of individual models reveals that—on average— the time of the early-to-late transition is significantly correlated with trial number (Fig 5C; p < 0.001). And despite its small absolute size, the average change in the time of transition predicted by the models was marginally significant when collapsed across taste (Fig 5D). The implication of these results is that the most “important” (i.e., well-studied) dynamical feature of GC ensemble taste responding changes slightly as the rat becomes accustomed to tasting.

Of course, it is possible that taste coding—the taste specificity of individual responses— could co-exist with unusual, “immature” response dynamics. To determine if the content coded by ensembles across dynamic states of taste responses changed across trials of novel taste exposure, we calculated how faithfully and distinctly the activity in the key states of the taste response coded the taste given to the animal. To calculate this, we performed Bayeisian decoding of taste identity, where for each recording, a gaussian classifier was trained on the ensemble responses corresponding to each state, labeled with both taste and state-number (i.e. if 4 tastes were given and each model contained 4 states, then the classifier would be trained to predict the probability that the test data came from each of the 16 possible states). Early and late taste-specific states were identified as the two states lasting at least 50ms, starting at least 50ms and before 2000ms post-stimulus in each trial, with the highest predicted probability of coding the correct taste that was administered in that trial. By assigning early-and-late states and decoding from them in this way, we were able to analyze the fidelity dynamic-states’ taste-coding while remaining agnostic to if the identity of the “early” and “late” states changed throughout a session.

Our examination of the relationship between trial number and taste-coding accuracy was modeled separately, using the same logistic function described previously, for the early and late-states (Fig 5E&H). Both the early and late states were significantly correlated with changes in the accuracy of taste-identity coding across trials in the session—that is, the logistic function fit significantly better than expected in each case (Fig 5F&I). This effect was of larger magnitude (and reached significance) in the late epoch (Fig 5J) than the early epoch (Fig 5H), with coding quality increasing by 30% (p < 0.001) across the session. Together, these results demonstrate that both the dynamics and coding content of GC taste responses evolve across the first 12 trials of a session, with the amount of information available in late-epoch activity notably increasing with experience.

### The rapid early changes in taste-processing largely reflect taste naivete

The fact that GC taste responsiveness stabilizes across the first dozen trials makes sense if one hypothesizes that taste processing is not completely innate but experience-dependent; given that rats are almost entirely taste naïve when the experiment begins (laboratory rats being raised with access to little other than bland chow), it could be said that the early trials are necessary to “tune up” the system. To test this possibility, we compared the above results to those obtained in identical analyses of a later session of taste exposure (Fig 6A&B).

**Fig. 6:**
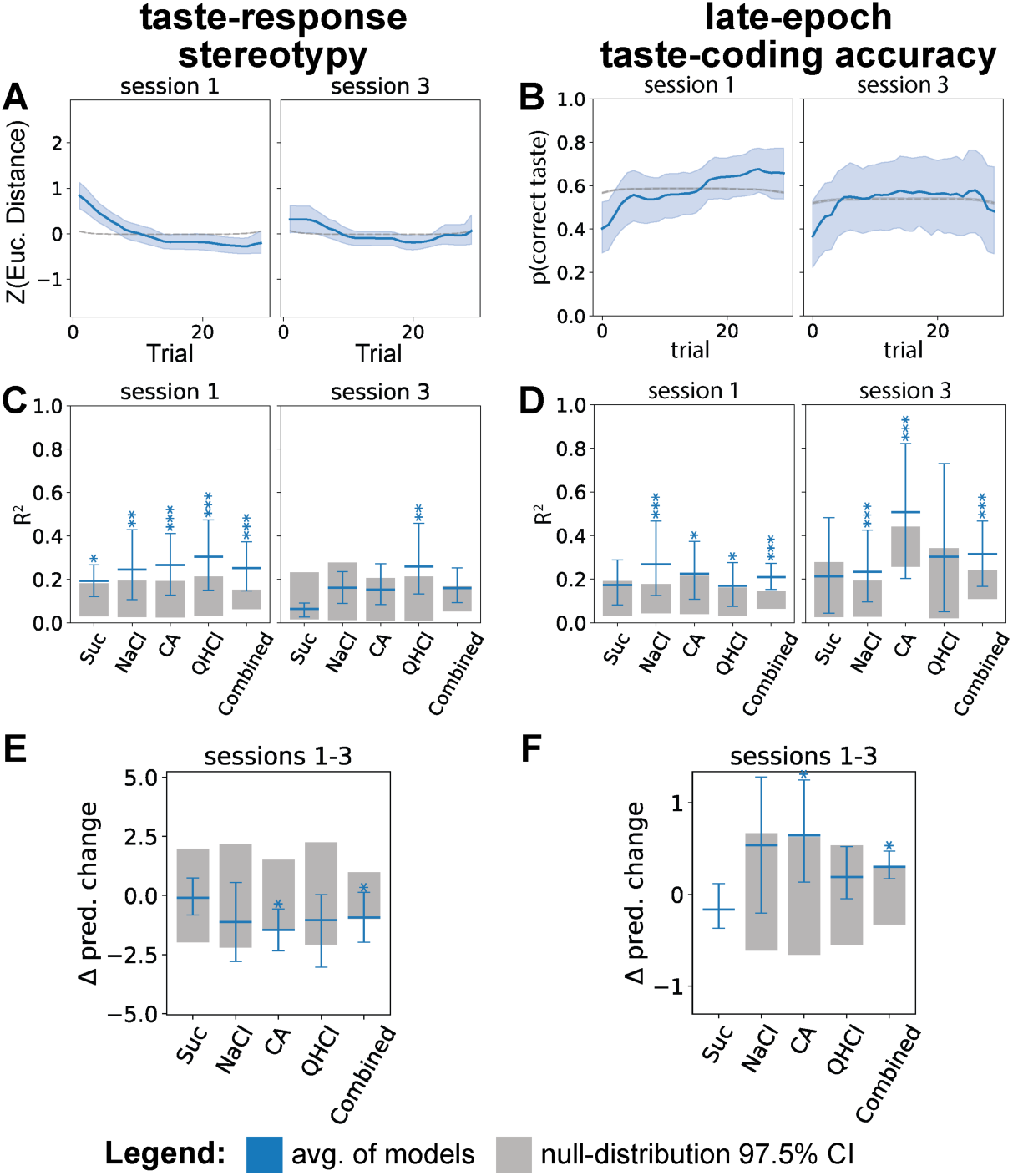
Rapid stabilization of GC taste responses reflects taste-naivete and taste novelty. **(A, B)** The 2 largest effects observed above—stereotypy of the taste response (A) and late-epoch identity-coding accuracy (B)— compared to the same effect in later (3^rd^) sessions for the same rats. See earlier figures for graphing conventions. There is almost no evidence of any across-session change in stereotypy in session 3 (A) and only a small, brief uptick in coding accuracy in session 3 (B). **(C, D)** Average coefficient of determination (R^2^) for the models in (A, B) reveals no increase in stereotype across session 3 (C), but the association of trial and coding accuracy in session 3 does achieve significance. **(E, F)** Direct comparison of sessions 1 & 3 reveals that GC taste responses exhibit significantly larger increases in stereotypy (E) and accuracy of taste-identity-coding (F) in session 1 than in session 3.

These results clearly supported our hypothesis. While stereotypy of responses in session 1 have a strong relationship to trial number, no such relationship was observed in the later session (Fig 6C), and the across-session change was significantly larger for session 1 than for session 3 (Fig 6D; only the citric acid response reached significance alone—almost assuredly a power issue—but the overall difference was reliable). And while we continued to observe a relationship between taste-identity-coding accuracy and trial number across the third session of taste responses (Fig 6B & E, see Discussion), the effect was considerably and significantly smaller than that observed in session 1 (Fig 6E; once again, the significance of the data was revealed in the across-taste analysis). Thus we conclude that the rapid changes observed in the first dozen taste responses in session one reflect an experience-dependent “tuning up” of the taste processing circuit occurring with novel taste exposures.

## Discussion

For decades, the corpus of taste research has operated under the assumption that taste-coding in GC is innate and stable, despite various lines of evidence suggesting that this is not entirely true. On the heels of the recent work demonstrating changes in GC-taste responses following novel-taste-exposure over days, this study aimed to investigate the extent to which GC taste responses are stable on the scale of individual trials, and determine if previously shown trial-to-trial variability in taste-responses can perhaps be mapped onto a familiarization-dependent phenomenon.

Initially, we observed distinct clusters of taste responses across trials within a session, indicating a clear separation between early- and late-session trial types. On the basis of the result that these analyses demonstrated taste-responses separated into two “regimes” of activity, we performed an analysis to identify the trial when this change happens, by identifying the bisection point where the average-response on either side of the bisection point best-summarize the responses across the sessions, and through this analysis confirmed that taste-responses indeed rapidly-evolve across the first dozen-or-so exposures to novel taste. It is important to note here, though, that these analyses are a representation of the composite of many sets of responses—one for each taste in each recording—and therefore represent an *average* shift in coding across trials. As a consequence, much like how averaging taste-responses over trials gives the appearance of smooth shifts in coding (Fig 2B), averaging over trials across recordings gives the appearance of a singular shift in coding across trials, when in reality it is more likely that there is animal-to-animal variability in the precise time at which taste responses enter into a stable regime of coding. The fact that sets of trials reliably form 2 clusters, but still exhibit significant overlap in which trials are binned into the “early-” or “late-session” clusters (Fig 3D), is highly suggestive that this is the case. Yet in the interest of focusing on other questions about these rapid changes, the question of variability in the time of this shift between sets of trials—whether that be taste or animal-specific—remains yet-to-be explored.

A key question we sought to answer was whether this shift in GC taste-coding was best described as a continuous, progressive change that converges upon a stable response, or if it is better-described as a sharp transition between two stable states. While—on the basis of the clustering results—it is tempting to believe the latter, it is possible for algorithms like hierarchical clustering to find clusters in data with “fuzzy borders” (Jain et al., 1999) that would characterize a progressively-changing response. Therefore, we quantified the within-group “stereotypy” of trials on either side of the “ideal” bisection point, to determine if these trials were bistable, which would preclude the idea of a converging change. Instead, we found that the earliest trials were significantly more varied around their template compared to the latter group, suggesting that this rapid change in taste responses did not represent a switch between stable states. Based on this finding, we went on to analyze our data under the assumption that this change across trials could be smooth, by modeling the data using a logistic-function. Over the average of many sets of trials, modeling of stereotypy across initial taste exposures is highly suggestive of a change resembling an exponential decay function that converges towards a “typical” response. The fact that—across sets-of-trials—these models consistently find that distance to the “average trial” decreases, indicates that “atypical” responses to novel taste are 1) a minority of responses in the first session, and 2) occur in the very beginning of the session, when taste is most novel. Altogether, these results are highly suggestive of a process where familiarization is occurring in real-time across individual exposures.

Beyond understanding that “something” about taste-responses is changing, we also wanted to know *what* about taste responses is changing. Therefore, we extracted the temporal dynamics of taste responses using Hidden-Markov-Modeling (HMM), which would allow us to formally disentangle changes in temporal-coding and population-coding. It was surprising to not find that the time of the early-to-late epoch transition does not consistently change with taste exposure, given that trial-to-trial variability in temporal dynamics of taste-responses is abundant. The fact that—for sucrose and NaCl responses—change in latency of the early-to-late epoch transition was strongly correlated with trial number, but that these models did not predict a significant, coherent change in the latency of this transition, suggests that the latency of this transition is likely progressively changing in opposite directions over trials across various sets of trials. Future analyses will need to examine whether the relationship between exposure and variability of temporal dynamics is better characterized as a change in the “noisiness” of the transition-time, rather than as a consistent change in the transition time. A key strength of this experiment was the use of multiple simultaneously presented tastes of varied palatability to assess familiarization. By simultaneously measuring the “decode-ability” of taste responses across trials, we found that the accuracy of taste-identity de-coding in the “late-epoch” of taste responses increases—a finding that is consistent with previous work from our lab identifying similar changes over days (Flores et al., 2022). Because classification accuracy depends on how distinctiveness of the data being classified, this finding is suggestive of the idea that taste-exposure “refines” taste-coding in GC during this epoch. Future analyses can make further use of our power to compare responses across classes and categories of taste by perhaps delving into the question of if responses to novel taste bear features of responses to aversive taste, or if they are their “own thing”.

Another key question that the presentation of multiple tastes allowed us to approach was whether only certain tastes elicited changes across familiarization—a question driven by well-described taste-specific differences in taste-neophobia (Reilly, 2018). We show somewhat mixed results. While all tastes exhibited changes across trials over some measure or another, sucrose was the only taste that did not exhibit a significant consistent change in stereotypy over trials, despite changes within individual trial-sets bearing significant correlation with trial-number. The relative stability of sucrose responses could reflect that rats may have had significant previous-life-experience consuming sweet carbohydrates, either via their chow, or from breastfeeding.

Furthermore, while the aggregate of tastes exhibited a significant change in stereotypy and accuracy over days—citric-acid was the only individual taste to exhibit significance in this measure as well, perhaps suggesting that citric acid elicits the strongest novelty-related effects. Despite these mixed results, the fact that almost every measure shown here demonstrates significant experience-dependent changes across the same sets of trials over the aggregate of responses to all the tastes, suggests that novel taste exposure of any kind drives changes that are common to all tastes.

Two central assumptions implicit in these experiments and analyses are that taste “experience” or familiarity is proportional to the amount of exposure, and that it is taste-specific. Therefore, the findings of “stereotypy” in this manuscript all represent averages of analyses performed within each “set” of trials: where each set contains each trial of responses to a specific taste in a recording—since there are 4 tastes, then each recording contains 4 “sets” of trials that are treated independently. While this was the simplest and most straight-forward way to categorize trials for these analyses, alternatives to these assumptions cannot be ignored. Namely, because these findings point to learning phenomenon, which are all subject to biophysically enforced time-constants underlying plasticity, it will be important to perform experiments that formally disentangle time from exposure-quantity. Furthermore, it is clear from abundant previous literature (Lubow, 2009; Flores et al., 2018, 2022; Wu et al., 2021) that the effect of taste-exposure is not truly taste-specific, but that the exposure to one novel taste changes how other tastes are perceived and processed in GC. While the experiments performed here de-couple time, exposure quantity, and specificity to some extent through random-ordering of taste exposures, the magnitude of this de-coupling is not large enough to show meaningful or interpretable differences in the results shown above when analyzed according to overall taste exposure number compared to taste-specific exposures.

Given that these results establish a reference nonlinear-temporal pattern for the rate at which taste-responses change, future experiments may choose to test the question of if exposure-quantity or time after exposure matters more. For example: one could attempt to “linearize” the change by carefully manipulating the timing of deliveries so early exposures have long separations in time, and each subsequent taste exposure happens in faster succession. To comprehensively answer the question of taste-specificity of experience, one could record taste-responses to novel tastes presented individually over many days.

Altogether, these results show compelling evidence that complements recent findings of familiarization-related changes in GC, and for the first time unveils the rapid nature of these changes. Although the evidence that GC taste responses change across familiarization is now abundant, it remains unclear if these changes are merely a phenomenon incidental to novel taste exposure, or if they could bear any behavioral relevance.

## Acknowledgements

This research was supported by US National Institutes of Health grants R01DC007703, T90DA032435, NSF/XSEDE computational resources allocation IBN180002, and the Brandeis National Committee—Los Angeles Chapter-endowed fellowship in Neuroscience and Biomedical Sciences. Thank you to Emma Barash, Thomas Gray, Roshan Nanu and Bradly T. Stone for assistance with various experiments and analyses.

## References

Arieli E, Younis N, Moran A (2022) Distinct Progressions of Neuronal Activity Changes Underlie the Formation and Consolidation of a Gustatory Associative Memory. J Neurosci 42:909–921.

Flores VL, Parmet T, Mukherjee N, Nelson S, Katz DB, Levitan D (2018) The role of the gustatory cortex in incidental experience-evoked enhancement of later taste learning. Learn Mem 25:587–600.

Flores VL, Tanner B, Katz DB, Lin J-Y (2022) Cortical taste processing evolves through benign taste exposures. Behav Neurosci 136:182–194.

Jain AK, Murty MN, Flynn PJ (1999) Data clustering: a review. ACM Comput Surv 31:264– 323.

Jones LM, Fontanini A, Sadacca BF, Miller P, Katz DB (2007) Natural stimuli evoke dynamic sequences of states in sensory cortical ensembles. Proc Natl Acad Sci 104:18772–18777.

La Camera G, Fontanini A, Mazzucato L (2019) Cortical computations via metastable activity. Curr Opin Neurobiol 58:37–45.

Lang L, Camera GL, Fontanini A (2023) Temporal progression along discrete coding states during decision-making in the mouse gustatory cortex. PLOS Comput Biol 19:e1010865.

Lubow RE (2009) Conditioned taste aversion and latent inhibition: A review. In: Conditioned taste aversion: Behavioral and neural processes, pp 37–57. New York, NY, US: Oxford University Press.

Matplotlib (2024) matplotlib.pyplot.imshow — Matplotlib 3.8.4 documentation. Available at: https://matplotlib.org/stable/api/_as_gen/matplotlib.pyplot.imshow.html [Accessed April 11, 2024].

Monk KJ, Rubin BD, Keene JC, Katz DB (2014) Licking microstructure reveals rapid attenuation of neophobia. Chem Senses 39:203–213.

Monti S, Tamayo P, Mesirov J, Golub T (2003) Consensus Clustering: A Resampling-Based Method for Class Discovery and Visualization of Gene Expression Microarray Data. Mach Learn 52:91–118.

Moran A, Katz DB (2014) Sensory Cortical Population Dynamics Uniquely Track Behavior across Learning and Extinction. J Neurosci 34:1248–1257.

Mukherjee N, Wachutka J, Katz DB (2019) Impact of precisely-timed inhibition of gustatory cortex on taste behavior depends on single-trial ensemble dynamics Colgin LL, Maffei A, Maffei A, Ghazanfar AA, Verhagen J, Mazzucato L, eds. eLife 8:e45968.

Nanu R, Svedberg D (2023) nubs01/blechpy. Available at: https://github.com/nubs01/blechpy [Accessed April 11, 2024].

Nanu RD, Murdy TJ, Levitan D, Scott S, Nelson SB, Katz DB (2021) Loss of Stk11 in basolateral amygdalar projection neurons impairs taste aversion learning by altering the temporal pattern of taste response plasticity in gustatory cortex. :2021.05.31.446460 Available at: https://www.biorxiv.org/content/10.1101/2021.05.31.446460v4 [Accessed April 9, 2024].

Reilly S (2018) 5 - Taste neophobia: Neural substrates and palatability. In: Food Neophobia (Reilly S, ed), pp 77–109 Woodhead Publishing Series in Food Science, Technology and Nutrition. Woodhead Publishing. Available at: https://www.sciencedirect.com/science/article/pii/B9780081019313000057 [Accessed April 9, 2024].

Sadacca BF, Mukherjee N, Vladusich T, Li JX, Katz DB, Miller P (2016) The Behavioral Relevance of Cortical Neural Ensemble Responses Emerges Suddenly. J Neurosci 36:655–669.

Schiff HC, Kogan JF, Isaac M, Czarnecki LA, Fontanini A, Maffei A (2023) Experience-dependent plasticity of gustatory insular cortex circuits and taste preferences. Sci Adv 9:eade6561.

Schoonover CE, Ohashi SN, Axel R, Fink AJP (2021) Representational drift in primary olfactory cortex. Nature 594:541–546.

scikit-learn (2022) sklearn.metrics.silhouette_score. Available at: https://scikit-learn/stable/modules/generated/sklearn.metrics.silhouette_score.html [Accessed April 11, 2024].

SciKit-Learn (2024) sklearn.naive_bayes.GaussianNB. Available at: https://scikit-learn/stable/modules/generated/sklearn.naive_bayes.GaussianNB.html [Accessed April 11, 2024].

SciPy (2022) Hierarchical clustering (scipy.cluster.hierarchy) — SciPy v1.13.0 Manual. Available at: https://docs.scipy.org/doc/scipy/reference/cluster.hierarchy.html [Accessed April 11, 2024].

SciPy (2024) SciPy documentation — SciPy v1.13.0 Manual. Available at: https://docs.scipy.org/doc/scipy/index.html [Accessed April 11, 2024].

Staszko SM, Boughter JD, Fletcher ML (2022) The impact of familiarity on cortical taste coding. Curr Biol CB 32:4914–4924.e4.

Vallat R (2024) pingouin.rm_anova — pingouin 0.5.4 documentation. Available at: https://pingouin-stats.org/build/html/generated/pingouin.rm_anova.html [Accessed April 11, 2024].

Wu C-H, Ramos R, Katz DB, Turrigiano GG (2021) Homeostatic synaptic scaling establishes the specificity of an associative memory. Curr Biol 31:2274–2285.e5.

